# Contrastive Functional Connectivity Defines Neurophysiology-informed Symptom Dimensions in Major Depression

**DOI:** 10.1101/2024.10.04.616707

**Authors:** Hao Zhu, Xiaoyu Tong, Nancy B. Carlisle, Hua Xie, Corey J. Keller, Desmond J. Oathes, Charles B. Nemeroff, Gregory A. Fonzo, Yu Zhang

**Affiliations:** Department of Bioengineering, Lehigh University, Bethlehem, PA, USA; Department of Psychology, Lehigh University, Bethlehem, PA, USA; Center for Neuroscience Research, Children’s National Hospital, Washington, DC, USA; Wu Tsai Neuroscience Institute, Stanford University, Stanford, CA, USA; Department of Psychiatry and Behavioral Sciences, Stanford University School of Medicine, Stanford, CA, USA; Veterans Affairs Palo Alto Healthcare System, and the Sierra Pacific Mental Illness, Research, Education, and Clinical Center (MIRECC), Palo Alto, CA, 94394, USA; Center for Brain Imaging and Stimulation, Department of Psychiatry, University of Pennsylvania Perelman School of Medicine, Philadelphia, PA, USA; Center for Neuromodulation in Depression and Stress, Department of Psychiatry, University of Pennsylvania Perelman School of Medicine, Philadelphia, PA, USA; Center for Psychedelic Research and Therapy, Department of Psychiatry and Behavioral Sciences, Dell Medical School, The University of Texas at Austin, Austin, TX, USA; Department of Electrical and Computer Engineering, Lehigh University, Bethlehem, PA, USA

## Abstract

**Background:** Major depressive disorder (MDD) is a prevalent psychiatric disorder characterized by substantial clinical and neurobiological heterogeneity. Conventional studies that solely focus on clinical symptoms or neuroimaging metrics often fail to capture the intricate relationship between these modalities, limiting their ability to disentangle the complexity in MDD. Moreover, patient neuroimaging data typically contains normal sources of variance shared with healthy controls, which can obscure disorder-specific variance and complicate the delineation of disease heterogeneity.

**Methods:** We employed contrastive principal component analysis to extract disorder-specific variations in fMRI-based resting-state functional connectivity (RSFC) by contrasting MDD patients (N=233) with age-matched healthy controls (N=285). We then applied sparse canonical correlation analysis to identify latent dimensions in the disorder variations by linking the extracted contrastive connectivity features to clinical symptoms in MDD patients.

**Results:** Two significant and generalizable dimensions linking distinct brain circuits and clinical profiles were discovered. The first dimension, associated with an apparent “internalizing-externalizing” symptom dimension, was characterized by self-connections within the visual network and also associated with choice reaction times of cognitive tasks. The second dimension, associated with personality facets such as extraversion and conscientiousness typically inversely associated with depression symptoms, is primarily driven by self-connections within the dorsal attention network. This “depression-protective personality” dimension is also associated with multiple cognitive task performances related to psychomotor slowing and cognitive control.

**Conclusions:** Our contrastive RSFC-based dimensional approach offers a new avenue to dissect clinical heterogeneity underlying MDD. By identifying two stable, neurophysiology-informed symptom dimensions in MDD patients, our findings may enhance disease mechanism insights and facilitate precision phenotyping, thus advancing the development of targeted therapeutics for precision mental health.

**Trial Registration:** Establishing Moderators and Biosignatures of Antidepressant Response for Clinical Care for Depression (EMBARC), NCT#01407094

## Introduction

Major depressive disorder (MDD) is a highly prevalent mental disorder with a lifetime prevalence of over 20% in women and 11% in men, impacting a substantial number of individuals globally. MDD is diagnosed based on the Diagnostic and Statistical Manual of Mental Disorders 5^th^ edition^1^. Unfortunately, such symptom-based diagnoses include patients with a bewildering mixture of symptoms and have resulted in substantial clinical and neurobiological heterogeneity, obscuring the underlying mechanisms of the cognitive and behavioral dysfunctions in MDD patients. This oversight may contribute to suboptimal treatment efficacy^2,3^ and hinder the development of more effective therapeutics, necessitating the dissection of MDD heterogeneity.

Prior studies have attempted to examine the clinical heterogeneity within MDD utilizing symptom-based dimensional approaches or subtyping analyses^4–6^. For instance, efforts have been made to identify symptom dimensions shared among patients and their associations with personality^7^, treatment outcomes^8^, or comorbid mental health conditions^9^. While current clinical measures are useful for inexpensively characterizing a patient symptom profile, they fall short in capturing underlying neurophysiological variations, limiting the discovery of neurobiological basis underlying the clinical heterogeneity. A recent systematic review^6^ reported that there is wide diversity of identified symptom dimensions and subtypes among previous studies, indicating the failure of addressing disease heterogeneity and the necessity of a more reliable and objective delineation of MDD pathological dimensions.

Neuroimaging techniques, such as functional magnetic resonance imaging (fMRI), have demonstrated promise in probing neurobiology of various psychiatric disorders. However, many previous neuroimaging studies have followed a case-control design that focuses on the group difference in aberrant brain circuits between MDD patients and healthy individuals, either during the resting state^10–12^ or in response to specific tasks^13^. Such an approach can only extract population-level information and has a limited ability to characterize the heterogeneity in MDD^14^. Recent research efforts have shifted to neuroimaging-based dimensional analyses to examine variations in brain structure and function beyond the conventional group-level comparison^15,16^. In particular, leveraging functional connectivity that measures neural coupling between brain regions, recent studies have revealed novel subtypes among mood and anxiety disorders^17,18^, which are not identifiable via a conventional case-control approach or clinically-defined MDD categories. To further enhance the clinical relevance of subtype or dimension discovery, increasing research efforts have delved deeper into not only dimensional patterns of brain circuits but also their behavioral or cognitive profiles^17,19–21^. Typically employing machine learning techniques such as canonical correlation analysis (CCA) or partial least squares, these studies have identified latent dimensions by jointly examining functional connectivity and symptom/behavioral measures in a data-driven manner^19–21^. These approaches revealed patterns of association between neurophysiological characteristics and specific symptom combinations, offering insights into the underlying mechanisms behind these disorders.

Neuroimaging data from patients contains both disorder-specific variation and disorder-irrelevant variation shared with healthy controls^22^. Given the high variability observed across individual-level brain metrics, one challenge in characterizing heterogeneity among clinical populations lies in the fact that it is usually muddied by overall normative variability. This has substantially decreased the signal-to-noise ratio and leads to suboptimal identification of meaningful patterns. In addressing this challenge, contrastive learning emerges as a powerful tool for training models to selectively retain group-specific features while filtering out confounding information inherent in the broader population. This process enables a focused analysis by isolating and emphasizing the distinctive characteristics pertinent to the target group. Encouragingly, some recent works have integrated the contrastive learning technique into the exploration of brain biomarkers in psychiatric disorders such as autism^22,23^, demonstrating its unique advantages in uncovering the intricate nuances of brain morphology^22^ and functional connectivity^23^ associated with psychiatric conditions. Contrastive learning holds great potential in capturing the inadequately explored pathology-relevant variations for defining symptom dimensions more precisely within MDD patients. Hence, we aim to shed light on the underlying heterogeneity specific to MDD through the identification of linked dimensions between clinical symptoms and MDD-specific neurophysiology components obtained by contrastive learning.

In this study, we applied a contrastive learning-based brain-symptom analytical framework^23^ to identify latent dimensions that link resting-state functional connectivity (RSFC) and clinical symptoms in MDD patients. This framework comprises two main steps. First, we conducted the contrastive principal component analysis (cPCA) on the fMRI-based RSFC data of MDD patients (N=233), with age-matched healthy controls (N=285) as background data, i.e. normative sources of variance to be removed from consideration in subtyping analyses. This process allowed us to extract contrastive RSFC features that disentangled disorder-specific variations from those shared with the healthy population. We then applied sparse CCA between these contrastive RSFC features and representative symptoms to reveal informative latent dimensions that link MDD-specific brain functional variations and clinical symptoms. Our study successfully identified two robust latent dimensions showing distinct neural circuit patterns and clinical profiles, primarily involving internalizing-externalizing symptoms and depression-protective personality. Rigorous stability analysis and cross-validation further confirmed their robustness and generalizability. We also examined associations of the identified dimensions with performance on various cognitive tasks, offering valuable insights into MDD-related cognitive dysfunction. Collectively, our contrastive RSFC-based dimensional approach provides a new avenue to examine neurophysiology-informed symptom dimensions for an improved understanding of mechanisms underlying MDD symptom heterogeneity. This may lead to a more objective classification of psychiatric conditions, thereby advancing targeted therapeutics for precision mental health.

## Materials and Methods

### Participants

#### Patient Population

The patient population used in our study include MDD patients from the *Establishing Moderators and Biosignatures of Antidepressant Response for Clinical Care for Depression* (EMBARC) dataset^24^. EMBARC is a large, randomized placebo-controlled clinical trial for examining biomarkers for MDD and antidepressant treatment response. Written informed consent was obtained from each participant under the Institutional Review Board (IRB) approved protocols at each of the four study sites, including University of Texas Southwestern Medical Center, Columbia University/Stony Brook, Massachusetts General Hospital, University of Michigan, University of Pittsburgh, and McLean Hospital. Subjects were required to meet the SCID criterion for an MDD episode and have a Quick Inventory of Depressive Symptomatology score^25^ of ≥14 at both screening and randomization visits. This dataset includes 296 MDD patients who were randomly prescribed sertraline or placebo for eight weeks, with the primary outcome of treatment response measured by the 17-item Hamilton Depression Rating Scale (HAMD_17_)^26^. For our brain-symptom linked dimension analysis, we included 25 baseline clinical scales and subscales: symptoms of depression (the 17-item HAMD; Quick Inventory of Depressive Symptomatology); Childhood Trauma Questionnaire subscales (emotional abuse, emotional neglect, physical abuse, physical neglect, sexual abuse)^27^; suicide-related symptoms (Concise Associated Symptoms Tracking^28^; Concise Health Risk Tracking-Self Report: propensity and risk scores^29^); Mood and Anxiety Symptoms Questionnaire subscales (anxious arousal, anhedonic depression, general distress)^30^; Mood Disorder Questionnaire^31^; personalities (NEO-Five Factor Inventory subscales, including neuroticism, extraversion, openness, agreeableness, and conscientiousness^32^; Standardised Assessment of Personality – Abbreviated Scale); Self-Administered Comorbidity Questionnaire^33^; Snaith-Hamilton Pleasure Score^34^; and Social Adjustment Scale overall mean score. Due to incomplete clinical measures, 63 subjects were excluded. The remaining 233 subjects (aged 18-65) were used for our analyses. Demographics of these subjects are listed in Supplementary Table S1.

#### Healthy Population

We compiled and harmonized 285 healthy controls (aged 18-77.5) as a background group for cPCA. Demographic information of these subjects is summarized in Supplementary Table S2. These subjects were selected by matching the age of the healthy population with the patient population (Kolmogorov–Smirnov test p = 0.052, Figure S1). Among them, 27 are from the EMBARC dataset and others are from the following three datasets. Specifically, we used 83 healthy controls from the University of California Los Angeles Consortium for Neuropsychiatric Phenomics (UCLA-CNP)^35^, which was approved by the IRB at University of California, Los Angeles and the Los Angeles County Department of Mental Health. 72 subjects were selected from the Amsterdam Open MRI Collection - Population Imaging of Psychology (AOMIC-PIOP1)^36^ dataset, which was approved by the faculty’s ethical committee at the University of Amsterdam. The remaining 103 subjects were from the Leipzig Study for Mind-Body-Emotion Interactions (LEMON)^37^ dataset, which was approved by the ethics committee at the medical faculty of the University of Leipzig.

### MRI Acquisition and Preprocessing

#### EMBARC

MRI data were acquired using 3T MRI systems at four different sites. At each site, resting-state fMRI data were scanned via T2* weighted images using a single-shot gradient echo-planar pulse sequence lasting for six minutes, with parameter settings: repetition time 2000 ms, echo time 28 ms, flip angle 90°, matrix size 64 × 64, voxel size 3.2 × 3.2 × 3.1 mm^3^, and 39 axial slices. Each subject underwent one or two runs of fMRI scans within a single day.

#### UCLA-CNP

MRI data were acquired on two 3T Siemens Trio scanners at UCLA. Resting-state MRI data were scanned using a T2*-weighted echoplanar imaging sequence lasting for 304 seconds, with the following parameters: repetition time 2000 ms, echo time 30 ms, flip angle 90°, matrix size 64 × 64, voxel size 3 × 3 × 4 mm^3^, and 34 slices.

#### AOMIC-PIOP1

MRI data were acquired on a Philips 3T scanner. During the resting state scans, participants were instructed to keep their gaze fixated on a fixation cross in the middle of the screen with a gray background and to let their thoughts run freely. Resting-state fMRI scanning lasted six minutes (i.e., 480 volumes with a repetition time 750 ms).

#### LEMON

MRI data were scanned using a 3T Siemens Verio scanner. Resting-state fMRI data were scanned using a T2_∗_-weighted gradient echo planar imaging sequence lasting for 15 minutes and 30 seconds. The sequence parameters were specified as follows: repetition time 1400 ms, echo time 30 ms, flip angle 69°, matrix size 88 × 88, voxel size 2.3 × 2.3 × 2.3 mm^3^, and 64 slices.

All the resting-state fMRI data were preprocessed using the fMRIPrep pipeline^38^ and aggregated into 100 regions-of-interest (ROIs) level time series according to the Schaefer parcellation^39^. RSFC features were then calculated as Pearson’s correlation coefficient in fMRI time series between every pair of ROIs. For MDD patients with multiple fMRI runs, we took the average of the RSFC across runs to generate the RSFC feature data used for the subsequent analysis. Given that the fMRI data from healthy controls were derived from different studies, we applied the well-established ComBat harmonization technique^40^ to their RSFC data to mitigate site effects. During the removal of site effects, age and gender were designated as biological covariates to be preserved, with EMBARC as the reference batch.

### Disentangling MDD-specific RSFC Variations via Contrastive PCA

In our study, we employed cPCA^41^ on the RSFC data of MDD patients as the target data and the RSFC data of healthy controls as the background data, aiming to pinpoint the disorder-specific foreground components by contrasting the healthy population. cPCA operates under the assumption that the target data comprises both domain-specific (foreground) information and domain-unrelated (background) variance. Utilizing the covariance matrix *C_p_* derived from the target data (MDD patients in our case) and the covariance matrix *C_bg_* derived from the background data (healthy controls), cPCA identifies the linear components most closely associated with the foreground data by subtracting the background covariance matrix from the target covariance matrix to obtain the foreground covariance matrix: *C_fg_* = *C_p_* − *αC_bg_*, where *α* represents a hyperparameter quantifying the degree of contrast. This hyperparameter was determined based on the cross-validation results through grid search. The top 200 contrastive principal components, which explain more than 70% of the data variance, were used in subsequent analyses. This selection was made because the dimensionality of informative components is constrained by the sample size of patients.

### Identifying Neurophysiology-informed Symptom Dimensions using sCCA

We first applied PCA to the clinical variables and retained 80% of the total variance to reduce dimensionality. We then applied sparse canonical correlation analysis (sCCA)^42^ to identify latent dimensions linking contrastive connectivity features with clinical symptoms. sCCA optimizes the correlation between two data matrices, thus yielding symptom dimensions maximally associated with neuroimaging features. Specifically, we utilized sCCA to analyze the disorder-specific components extracted from cPCA and corresponding clinical measures, aiming to identify neurophysiology-informed symptom dimensions. Sparsity constraint is imposed on both connectivity features and clinical measures to improve interpretability and alleviate overfitting. The overall minimization objective is formulated as −*Cov*(*Au*, *Bv*) + *λ*_1_|*u*|_1_ + *λ*_2_|*v*|_1_, where *A* and *B* are data matrices of connectivity features and clinical measures respectively, *u*, *v* are the FC and symptom dimension loadings, and *λ*_1_, *λ*_2_ are sparsity hyperparameters on the loadings. We performed grid search to find the hyperparameters with best performance.

### Cross-Validation and Statistical Significance Verification

We performed 10 repetitions of 10-fold cross-validation to evaluate the generalizability and stability of identified latent dimensions. In each fold, 90% of the patients and all healthy subjects were compiled as the training set of cPCA. As the order of identified CCA dimensions might vary across folds due to training data variance, we employed the dimensions identified from the whole dataset as reference dimensions to align the dimensions acquired from different folds. Pearson’s correlation coefficient was then computed to assess the generalizability of the identified dimensions on the validation set. Intra-class correlation coefficient was employed to confirm the stability of dimension loadings across cross-validation folds.

Furthermore, to evaluate the significance of the identified dimensions, we conducted permutation tests 1000 times. Specifically, we randomly permuted the clinical symptoms for each patient to others and followed the same procedures to generate the Pearson correlation R values on cross validation test set as described above. Test R values from dimensions with the same ranking across different folds were aggregated to generate the null distribution.

### Network-level Connectivity Importance

Network-level connectivity provides a broader view of which neural systems and their interactions contribute most to the identified latent dimensions. To measure this, we first retained only the top 10% loadings with the highest absolute values, setting the rest to zero to exclude the less important and stable connections. Then, we averaged all absolute loadings within and between each pair of networks to obtain the network-level connectivity importance.

### Post-hoc Examination of Associations between Dimensions and Cognitive Task Performance

We investigated the correlation between the FC scores of identified dimensions and the cognitive task performance of MDD patients. This analysis may provide insights into how different cognitive abilities, such as psychomotor slowing^43^, cognitive control^44^, working memory^45^, reward learning^46^, and resolution and adjustment behavior in response to emotional conflict^47^, relate to these brain-symptom latent dimensions. These cognitive and emotional measurements are selected because they are potential predictors for MDD treatment response^24^, thus having the probability to be correlated with the identified MDD-specific dimensions. The specific task item we included can be found in Table S3.

## Results

### Contrastive RSFC Defines Two Generalizable Brain-Symptom Linked Dimensions in MDD

We applied the contrastive learning-based brain-symptom association identification framework to the 233 MDD patients. Two dimensions with generalizable association between neurophysiology and symptom profile have been identified through 10 rounds of 10-fold cross-validation and permutation test. The composition and essence of the first identified dimension was interpreted, based upon loadings, to indicate an axis of internalizing-externalizing symptoms (R_train_ = 0.551, R_cv_ = 0.287, p_permutation_ < 0.001, Figures 2, S4). A higher internalizing-externalizing symptoms dimension score indicates a greater tendency towards internalizing behavior such as lack of pleasure or energy, and a lower tendency towards externalizing behavior such mania and anger attack. The second identified dimension incorporates the concept of depression-protective personality (R_train_ = 0.437, R_cv_ = 0.207, p_permutation_ = 0.001, Figure 2, S4). Individuals with higher scores on this personality dimension score exhibited higher scores on personality facets typically negatively associated with depression symptoms, such as extraversion and conscientiousness, while a low dimension score indicates higher scores on personality facets typically associated with greater depression symptoms, such as neuroticism^48^. Both dimensions significantly outperformed the performance yielded from its non-contrastive counterpart integrating standard PCA and sCCA (paired t-test: internalizing-externalizing symptoms dimension: p=5.2×10^-4^; depression-protective personality dimension: p = 0.008), demonstrating the unique advantage of contrastive learning for improving the identification of brain-symptom associations. Afterward, we calculated the intra-class correlation coefficient (ICC) of dimension composition in each cross-validation fold to assess the stability of the dimensions. These two dimensions both showed high stability. For internalizing-externalizing symptoms, the ICC value is 0.87 (95% CI: [0.87, 0.88]) for FC loadings, 0.92 (95% CI: [0.88, 0.96]) for symptom loadings. For depression-protective personality, the ICC value is 0.94 (95% CI: [0.94, 0.94]) for FC loadings, 0.94 (95% CI: [0.90, 0.97]) for symptom loadings (Figure S4).

**Figure 1.**
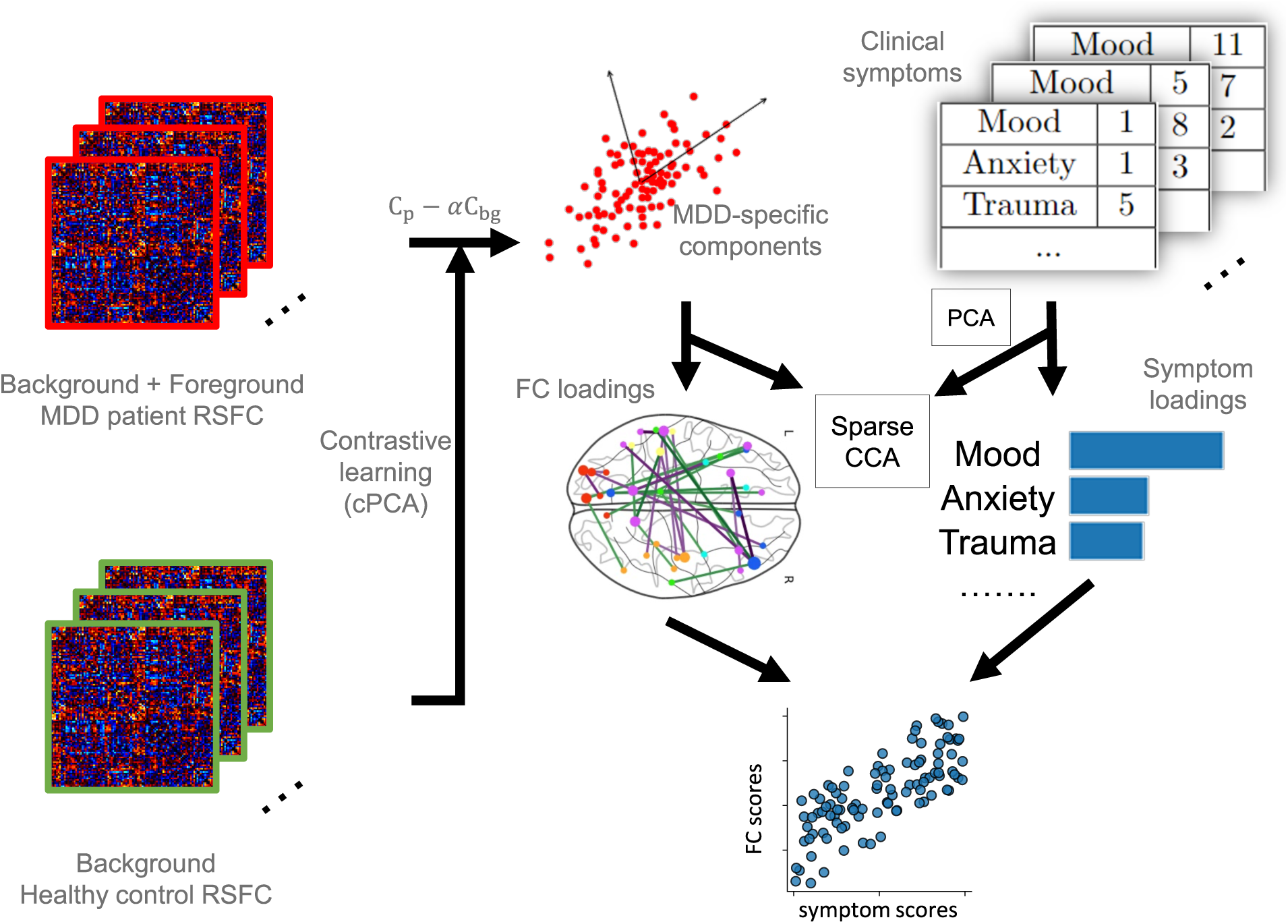
Illustration of the proposed framework. MDD patient RSFC data is considered to include both disorder-specific information (foreground) and shared noise with healthy control data (background). Contrastive learning is used for removing the background noise from patient data to generate MDD-specific components. Specifically, the covariance matrix of patient RSFC (*C_p_*) and healthy control RSFC (*C_bg_*) are computed to represent the group variances. MDD-specific components are derived from the updated covariance matrix *C_p_* − *αC_bg_*, which represents the foreground variance, and *α* represents a hyperparameter quantifying the degree of contrast. These components are used as MDD-specific RSFC features in the next step. Subsequently, sparse CCA is performed to identify the linked dimensions between neurophysiology and PCA-transformed symptoms, represented by FC and symptom loadings. These loadings serve as linear combinations which project the RSFC features and symptoms into a common space with high correlations.

**Figure 2.**
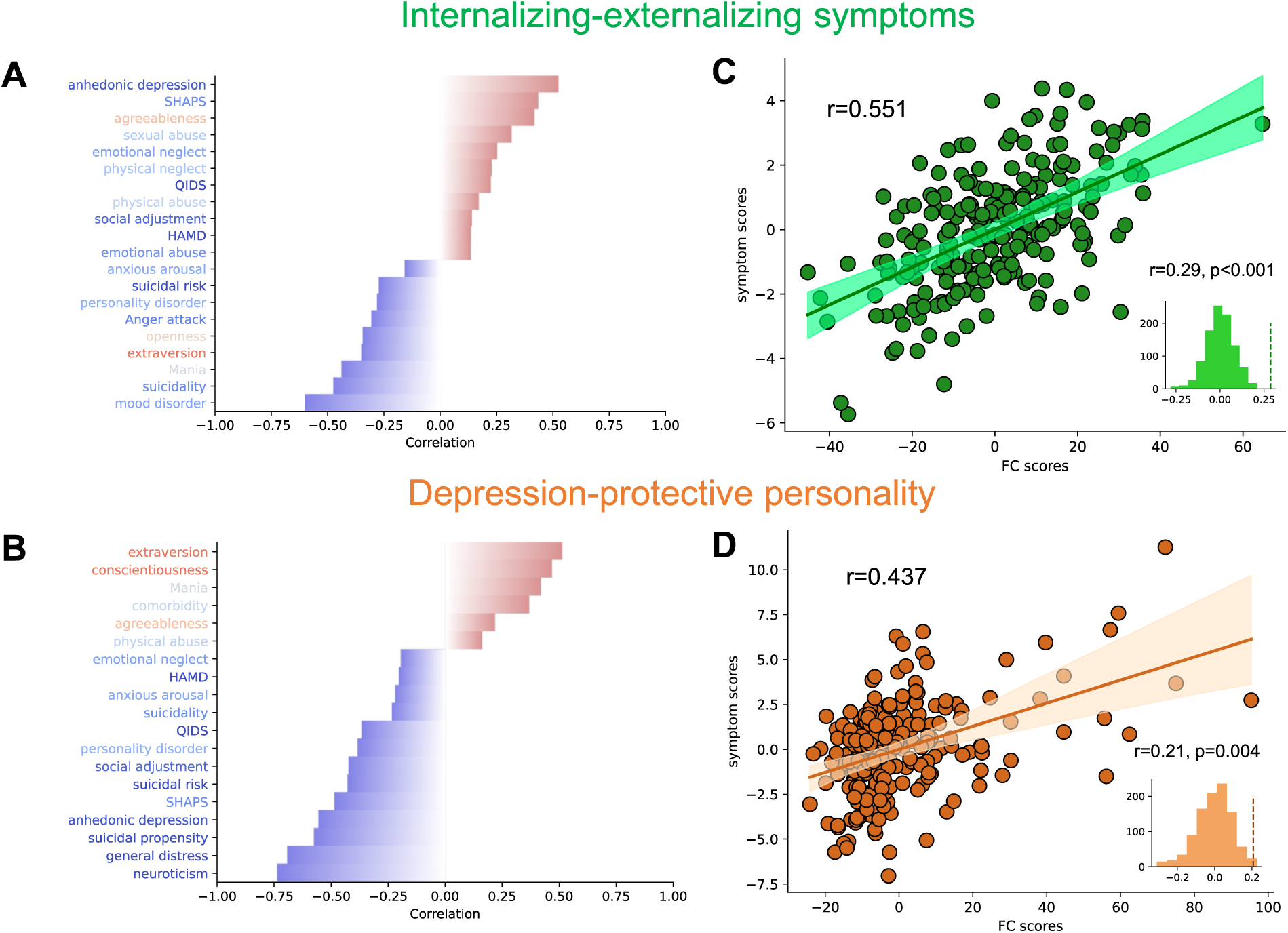
Identified linked dimensions between fMRI RSFC features and clinical measurements. (A) Top correlations between the symptom score of internalizing-externalizing symptoms dimension and 25 clinical measures. Leading negative correlations include Mood Disorder Questionnaire, Concise Associated Symptoms Tracking, and Altman Self-Rating Mania Scale. Anhedonic depression of Mood and Anxiety Symptoms Questionnaire is the leading positive correlation. The text color of the clinical measures represents the direction of the differences between healthy controls and MDD patients. Red indicates that healthy controls have higher average scores, while blue indicates that patients have higher average scores. (B) Top correlations of depression-protective personality dimension. NEO-Five Factor Inventory subscales (extraversion, conscientiousness, neuroticism) contribute the most. (C, D) Correlation between FC and symptom scores of the two linked dimensions on all patients. Cross-validation permutation results of 1000 repeats are shown in the small panels.

### RSFC Signatures Associated with MDD Symptoms

Next, we investigated the most important ROI-level and network-level connectivity within each dimension. (Figure 3). In the internalizing-externalizing symptoms dimension, the correlation between the left and right cuneus and other regions, particularly the left fusiform gyrus, the left inferior temporal gyrus, and the right fusiform gyrus, contributes the most. In terms of anti-correlation, the connection between the left and right posterior cingulate cortex and other regions, mainly the left superior occipital gyrus and the right middle occipital gyrus, contributes the most. On the network level, the dimension is characterized by the self-connection of visual network, the connection between limbic network and sensorimotor network, and the connection between limbic network and dorsal attention network. In depression-protective personality dimension, the correlation between the right middle cingulate cortex and the left superior parietal cortex contributes the most. In terms of anti-correlation, the key connections are between the left fusiform gyrus, the left inferior temporal gyrus, and the left superior parietal cortex, as well as with the left inferior frontal gyrus and the left middle frontal gyrus. On network level, this dimension is highly related to self-connections within the dorsal attention network.

**Figure 3.**
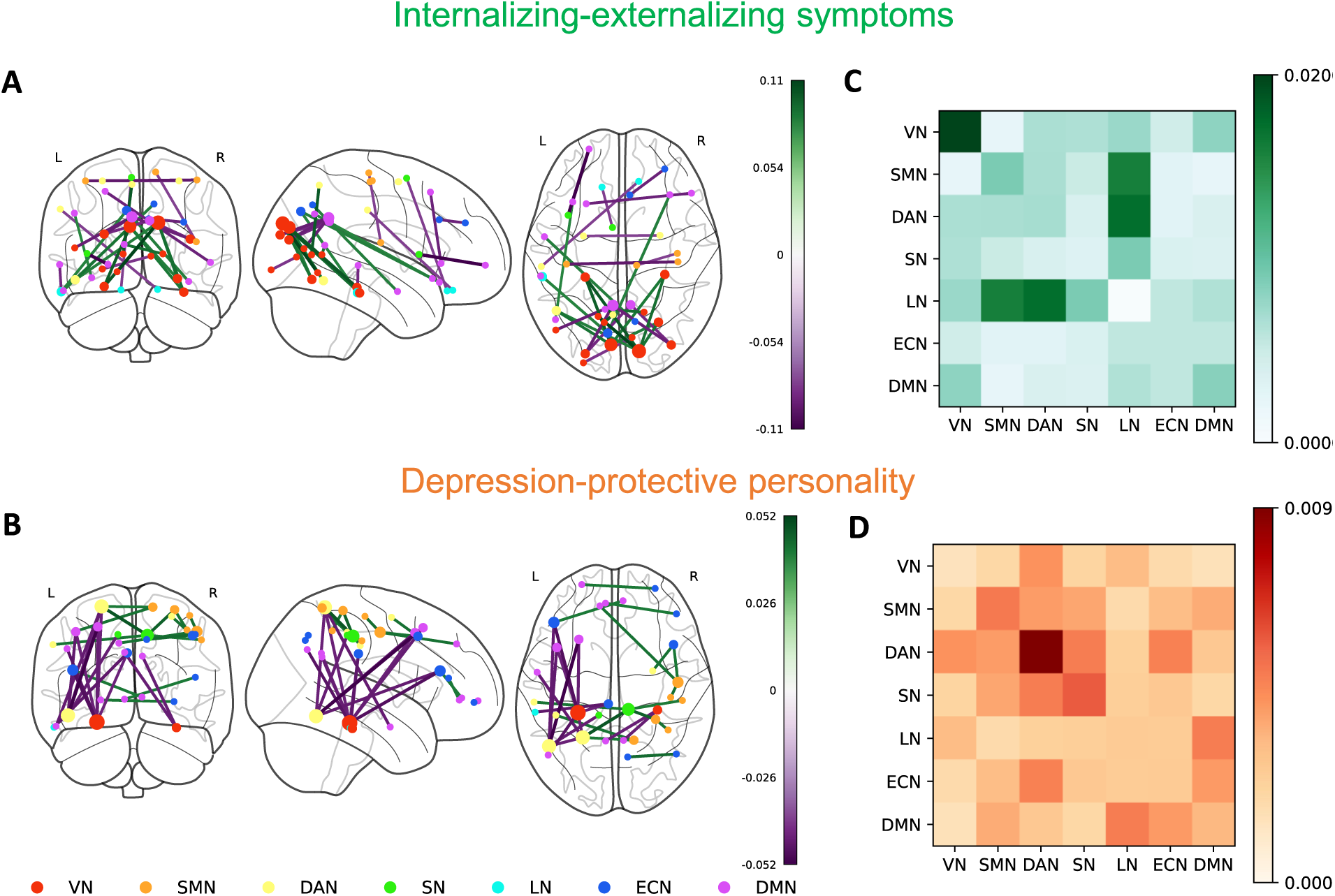
RSFC loadings and network-level importance of internalizing-externalizing symptoms dimension (A,C) and depression-protective personality dimension (B,D). (A, B) Top 15 positive and negative loadings of FC dimensions. Node size indicates the involvement of ROI in top loadings. (C, D) Network-level importance within two FC dimensions among 7 brain networks: visual network, sensorimotor network, dorsal attention network, salient network, limbic network, executive control network and default mode network. The values of network-level importance are calculated by first keeping only the top 10% loadings with highest absolute values and setting the rest to 0, and averaging all absolute loadings between every pair of networks.

### Dimensions Are Associated with Cognitive Task Performance

Internalizing-externalizing symptoms score showed a significant correlation with reaction time of choice reaction time task (r=0.20, p_fdr_ =0.0084) and reaction time difference in a flanker task (r=0.18, p_fdr_ =0.024). This indicates that a higher score on this dimension is associated with longer reaction time, suggesting lower performance. Reaction time difference in the flanker task measures selective attention and inhibitory function; longer reaction times indicate a greater interference effect from incongruent stimuli. Depression-protective personality dimension score had significant correlations with multiple cognitive tasks, including the accuracy of A-not-B task (r=-0.24, p_fdr_=0.0016), reaction time of choice reaction time task (r=0.31, p_fdr_ =2.8×10^−5^), and both the accuracy difference (r=-0.15, p_fdr_ =0.046) and reaction time difference (r=0.20, p_fdr_ =0.0084) in the flanker task. This suggests that this dimension has a wider range of correlations with different cognitive tasks measuring working memory, interference adjustments, and cognitive control. No significant correlations were observed between these two dimensions and the word fluency task, emotion conflict task, or probabilistic reward task (Figure 4).

**Figure 4.**
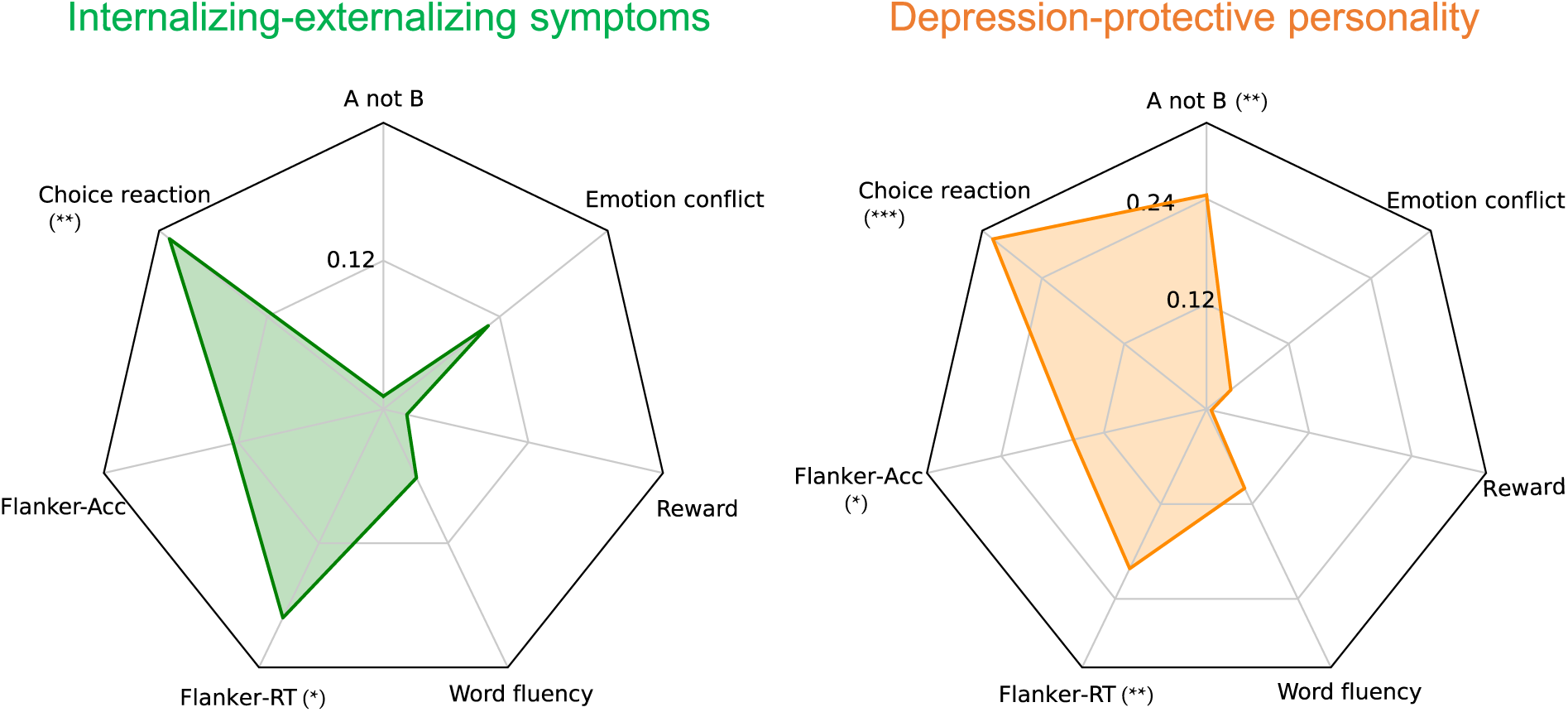
Pearson correlation between two FC dimensions and cognitive task performances. Left: internalizing-externalizing symptoms dimension; Right: depression-protective personality dimension. Cognitive task items include: 1) the overall accuracy of the A not B task, 2) the overall reaction time of the choice reaction time task, 3) the accuracy difference between incongruent and congruent trials in the flanker task, 4) the reaction time difference between incongruent and congruent trials in the flanker task, 5) the total valid number of words in the word fluency task, 6) the overall response bias in the probabilistic reward task, and 7) the reaction time difference between incongruent-following and congruent-following incongruent trials in the emotion conflict task. *** p_fdr_<0.001, **p_fdr_<0.01, * p_fdr_<0.05.

We also examined the capabilities of these dimensions in predicting antidepressant treatment response using chi-square test. However, no significant performance was observed for either dimension in differentiating remission versus non-remission based on whether the FC dimension scores were above or below the median (internalizing-externalizing symptoms dimension: p=0.27 for sertraline arm, p=1.00 for placebo arm; depression-protective personality dimension: p=0.40 for sertraline arm, p=0.49 for placebo arm) (Supplementary Table S4).

## Discussion

In this study, we implemented a data-driven framework that integrated contrastive machine learning with multivariate correlation analysis to uncover MDD-specific brain functional variations and their associations with clinical symptoms. We identified two robust and generalizable latent dimensions linking neurophysiological and clinical profiles, offering novel and objective biomarkers for dissecting heterogeneity in MDD.

While neuroimaging data conveys important information about circuit dysfunction in MDD patients, it also contains undesired variance shared with healthy individuals. Conventional dimensional methods obtain and identify behavior or symptom-related biomarkers from clinical populations, which may result in suboptimal findings without removing the variance shared by healthy populations. Alternatively, contrastive learning-based frameworks can better extract disorder-specific neurophysiology components through data distillation. In our results, the dimensions derived from contrastive learning show a reduced correlation with age compared to those without contrastive learning (Figures S2, S3). This suggests that contrastive learning successfully removed some disorder-unrelated components, such as age. Other than the undesired variance, MDD itself also exhibits substantial neurobiological and clinical heterogeneity. Previous studies based on case-control analysis or dimensional approaches considering only clinical symptoms or neuroimaging may therefore inadequately address the heterogeneity^6,49^. Our framework links both modalities, addresses the underlying heterogeneity, and provides interpretability.

For the two neurophysiology-informed symptom dimensions we have identified, internalizing-externalizing symptoms dimension is associated with key brain regions such as the cuneus and fusiform gyrus, consistent with findings from studies on neurodevelopmental trajectories of internalizing-externalizing symptoms^50^. Our results suggest that these associations likely persist into adulthood. Our network-level analysis on this dimension aligns with the literature, which identifies the visual network and dorsal attention network as strong predictors of internalizing-externalizing traits^51^. Our analysis of the behavioral task is also consistent with findings in the literature that associate cuneus activation with facilitating vigilance^52^.

The depression-protective personality dimension is marked by personalities such as extraversion, conscientiousness and neuroticism. Studies have identified a strong association between key brain regions, such as the fusiform gyrus, and personality disorders^53,54^, which is consistent with our findings. A recent study also identified the significant correlations between dorsal attention network and personalities including neuroticism and agreeableness^55^, and borderline personality disorder^56^. Additionally, dorsal attention network has also shown great importance in multiple cognitive control tasks^57,58^. All these results align with our findings to the depression-protective personality dimension.

Despite our contributions, this study has several limitations. First, while cPCA is effective for extracting disorder-specific components, it requires fine-tuning of hyperparameters to control the degree of contrast with the background data (i.e., healthy population). Future research could focus on developing automatic frameworks for extracting contrastive connectivity features. Additionally, our study harmonized data from different sources to enlarge the sample size, enhancing the robustness of the identified latent dimensions, followed by verifying their generalizability through rigorous cross-validation. However, future work should further involve independent datasets with comparable clinical measurements to replicate our findings. Lastly, given the recognized importance of subcortical regions and cerebellum in depression^59,60^, extending the analysis to these regions may provide a more comprehensive understanding of brain dysfunctions associated with MDD psychopathology.

In summary, this study employed a combination of contrastive learning and sparse canonical component analysis on a depression patient dataset, unveiling two generalizable disorder-specific dimensions linking neurophysiology and symptom profiles. Our findings hold potential for advancing the understanding of disease mechanisms, facilitating precise diagnosis, formulating individualized treatment plans, and fostering the development of innovative treatment methods.

## Acknowledgements

This work was supported by NIH grant nos. R01MH129694, R21MH130956, R21AG080425, Alzheimer’s Association Grant (AARG-22-972541), and Lehigh University FIG (FIGAWD35), CORE, and Accelerator grants. Portions of this research were conducted on Lehigh University’s Research Computing infrastructure partially supported by NSF Award 2019035. G.A.F. was also supported by philanthropic funding and NIH grant nos. R01MH132784 and R01MH125886, and grants from the One Mind – Baszucki Brain Research Fund, the SEAL Future Foundation, and the Brain and Behavior Research Foundation.

## Supplementary Materials

**Table S1.**
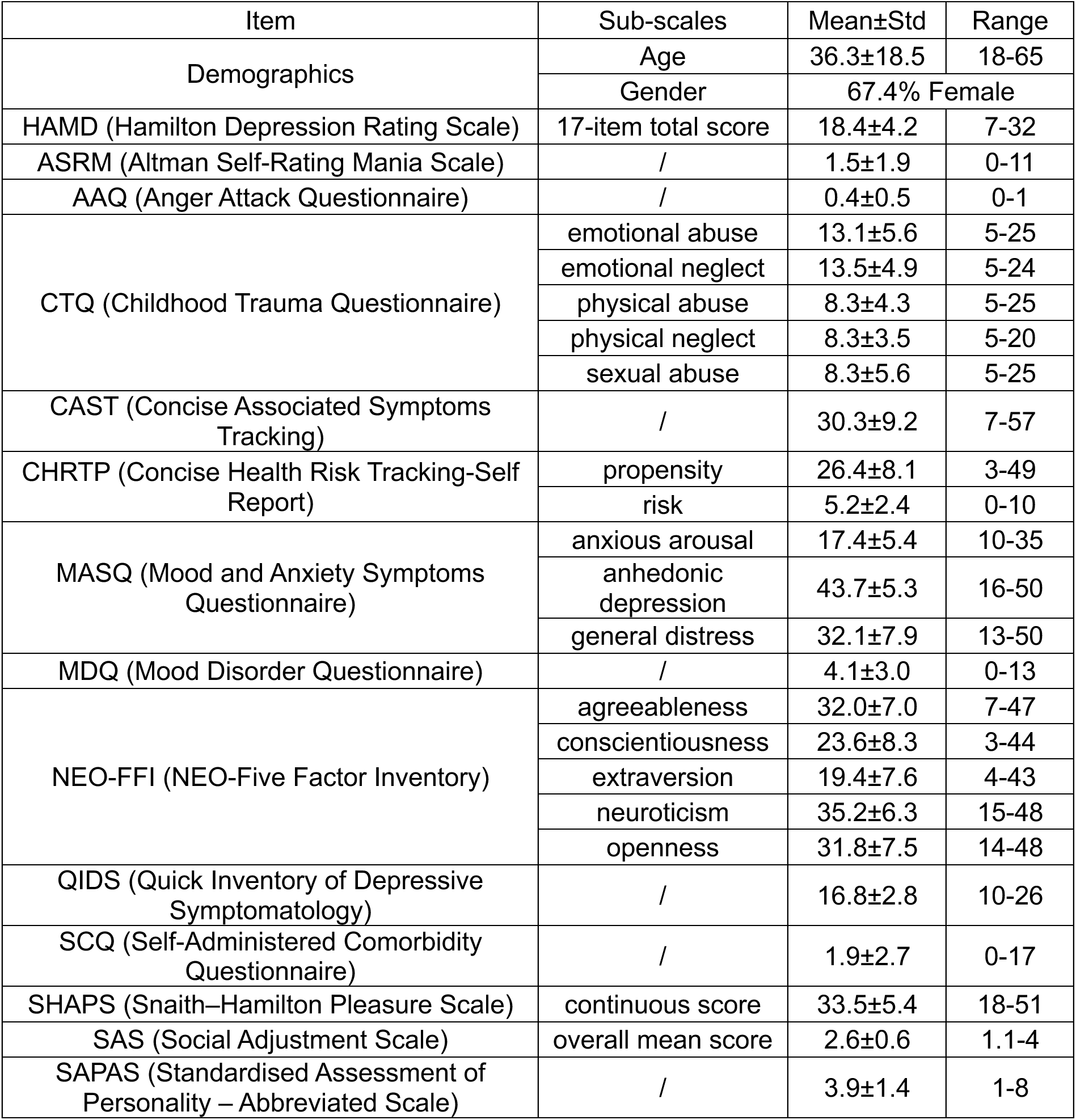
Distribution of demographic information and clinical measurements of EMBARC MDD patients included in our study (N=233)

**Table S2.** Demographic information of healthy controls included in our study.

|  | UCLA CNP<br>(N=83) | LEMON<br>(N=103) | AOMIC<br>(N=72) | EMBARC<br>(N=27) | Total<br>(N=285) |
| --- | --- | --- | --- | --- | --- |
| Age | 35.31±8.26 | 45.22±20.72 | 21.96±1.86 | 42.75±12.84 | 36.73±16.48 |
| Female % | 43.3 | 33.0 | 56.9 | 60.8 | 46 |

**Table S3.** Cognitive and emotional tasks included in our study and their measurements.

| **Task** | **Item** | **Measurements** |
| --- | --- | --- |
| A not B task | Overall accuracy | Working memory |
| Choice reaction time task | Overall reaction time | Interference and post-error adjustments |
| Flanker task | Accuracy difference between incongruent and congruent trials | Cognitive control, conflict monitoring |
|  | Reaction time difference between incongruent and congruent trials | Cognitive flexibility, conflict processing speed |
| Word fluency task | Total valid number of words | Psychomotor slowing |
| Probabilistic reward task | Overall response bias | Reward learning |
| Emotion conflict task | Reaction time difference between incongruent-following and congruent-following incongruent trials | Resolution and adjustment behavior in response to emotional conflict |

**Table S4.** Chi-square test between the population distribution in high/low FC dimension score groups and remission rates within different treatment groups (sertraline and placebo). Remission is defined as HAM-D score ≤ 7 at week 8. For each dimension, high/low groups are divided by the median FC dimension score from all population. All p-values > 0.05, indicating that there is no significant relation between high/low dimension scores and remission.

| Internalizing-externalizing symptoms dimension |  |  |  |  |  |  |
| --- | --- | --- | --- | --- | --- | --- |
| Sertraline | Observed |  | Expected |  | Statistics |  |
| | Remission | Non-remission | Remission | Non-remission | $\chi^2$ | 1.20 |
| High group | 14 | 42 | 17.19 | 38.81 | p | 0.27 |
| Low group | 21 | 37 | 17.81 | 40.19 | samples | 114 |
| Placebo | Observed |  | Expected |  | Statistics |  |
| | Remission | Non-remission | Remission | Non-remission | $\chi^2$ | 0.00 |
| High group | 16 | 45 | 15.89 | 45.11 | p | 1.00 |
| Low group | 15 | 43 | 15.11 | 42.89 | samples | 119 |
| Depression-protective personality dimension |  |  |  |  |  |  |
| Sertraline | Observed |  | Expected |  | Statistics |  |
| | Remission | Non-remission | Remission | Non-remission | $\chi^2$ | 0.71 |
| High group | 21 | 39 | 18.42 | 41.58 | p | 0.40 |
| Low group | 14 | 40 | 16.58 | 37.42 | samples | 114 |
| Placebo | Observed |  | Expected |  | Statistics |  |
| | Remission | Non-remission | Remission | Non-remission | $\chi^2$ | 0.48 |
| High group | 17 | 40 | 14.85 | 42.15 | p | 0.49 |
| Low group | 14 | 48 | 16.15 | 45.85 | samples | 119 |

**Figure S1.**
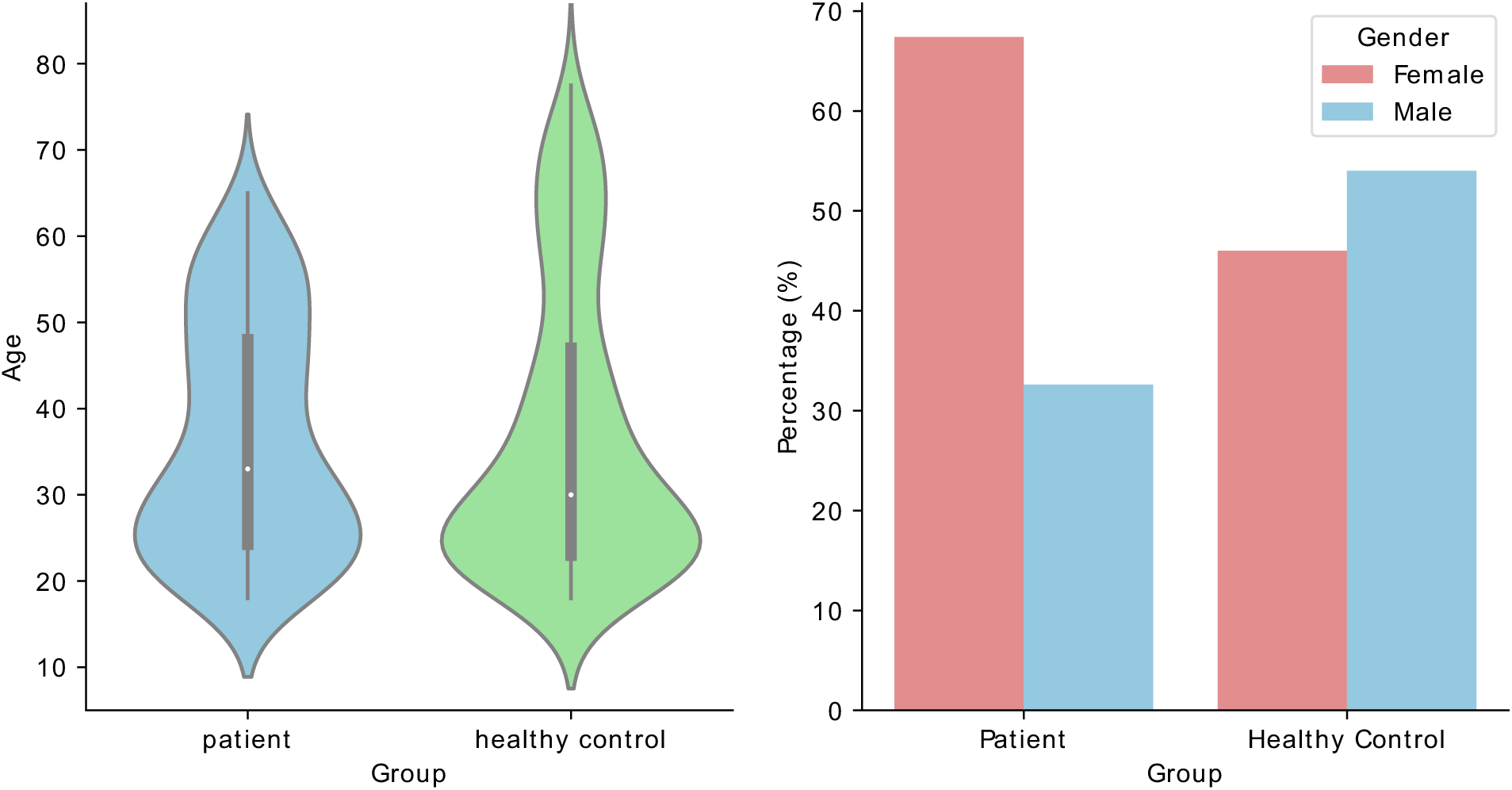
Demographic information comparison between patient (N=233) and healthy control groups (N=285). The age distribution of the patient population and the healthy population was matched, with a Kolmogorov-Smirnov test yielding a p-value of 0.052.

**Figure S2.**
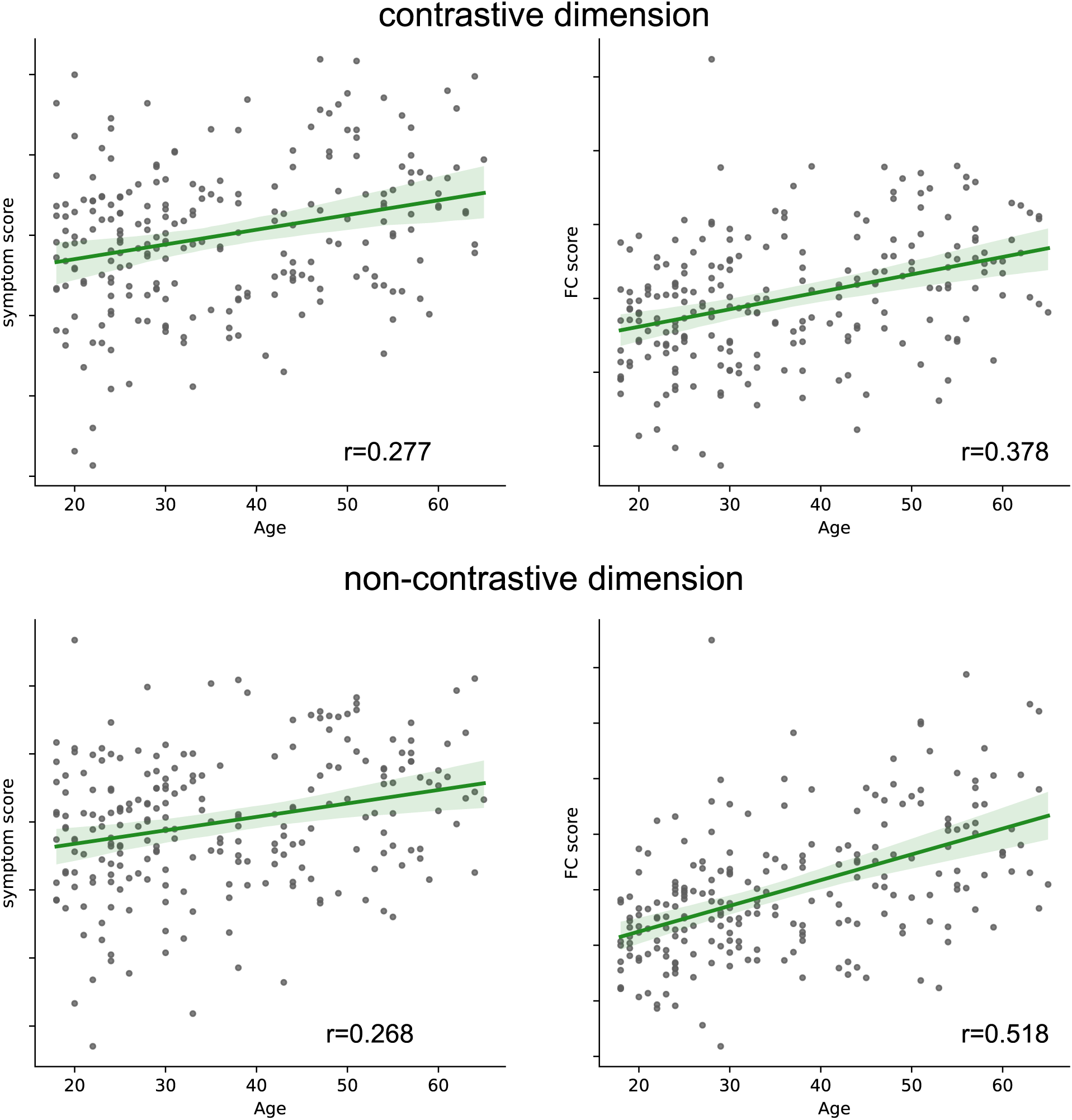
Comparison of the age-correlation of the best symptom/FC dimensions derived from cPCA and regular PCA components. Contrastive learning marginally reduced the age-related component in its FC dimension, with a Fisher’s z test showing near-significant results (p = 0.059).

**Figure S3.**
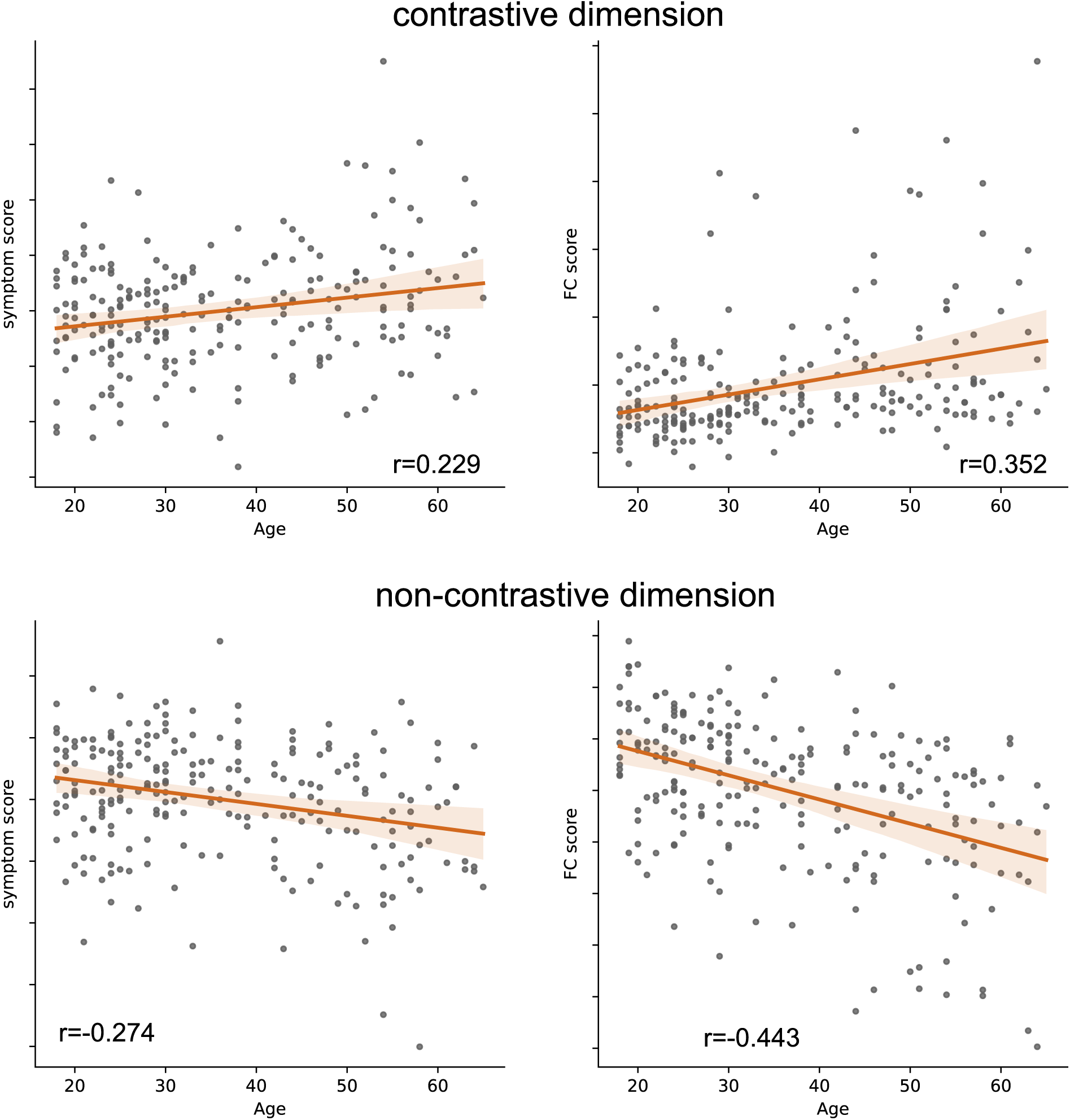
Comparison of the age correlation in the second-best symptom/FC dimensions derived from cPCA and regular PCA components. Contrastive learning reduced the age-related component in both symptom and FC dimensions, but the reduction was not statistically significant (p > 0.1).

**Figure S4.**
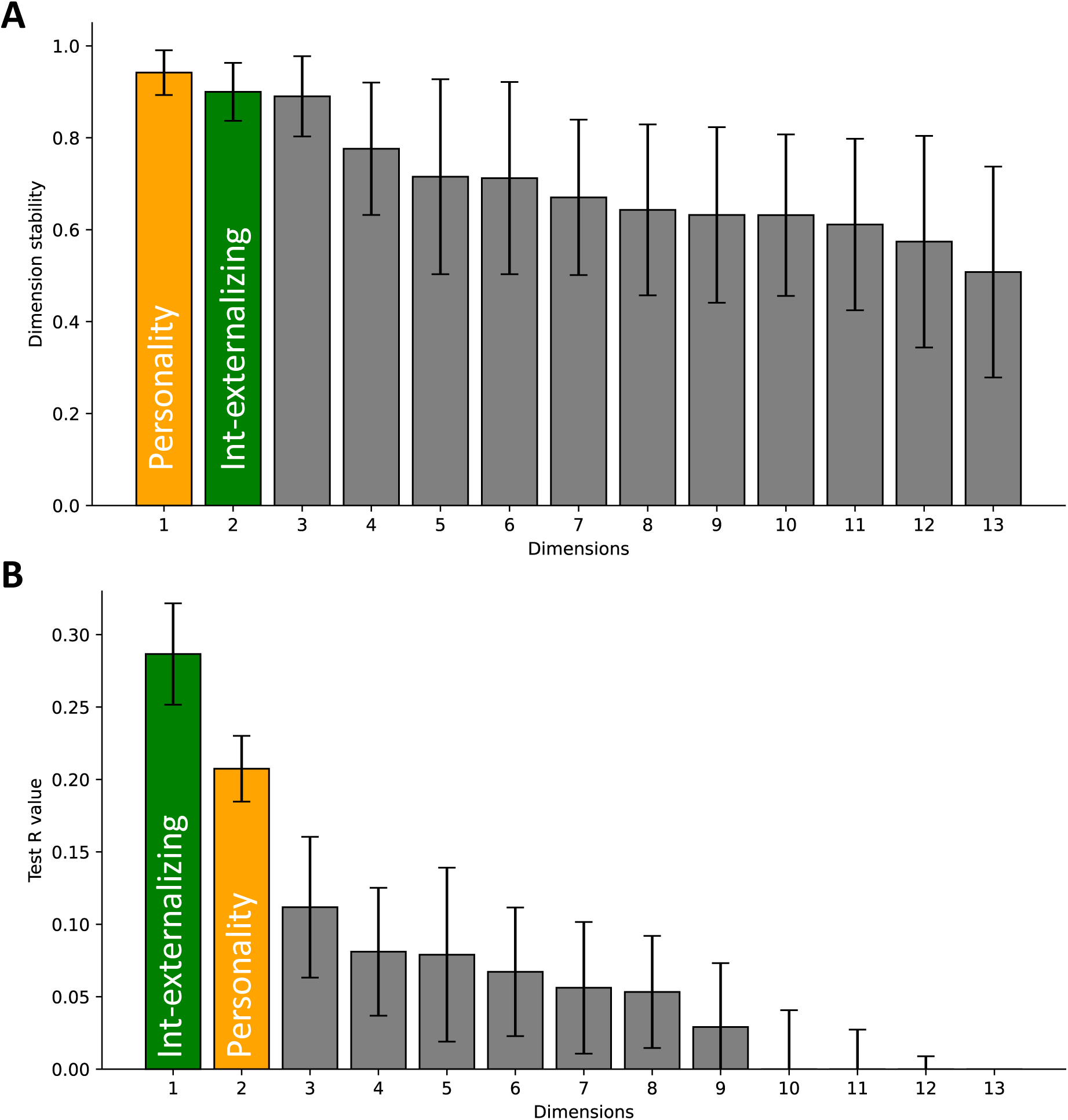
(A) Overall dimension stability for all 13 dimensions given by sCCA, measured by the average of the cosine similarity between each pair of FC dimensions and each pair of symptom dimensions from 10×10 folds. All dimensions have an overall stability over 0.5. (B) For the 13 dimensions, only two dimensions survived FDR correction in permutation test, shown in colors. The raw p-values are: p<0.001 for internalizing-externalizing symptoms dimension, and p=0.001 for depression-protective personality dimension.

